# MEK1/2 in Rhabdomyosarcoma

**DOI:** 10.1101/629766

**Authors:** Kenneth A. Crawford, Megan M. Cleary, Cora A. Ricker, Matthew N. Svalina, John F. Shern, Hung-I Harry Chen, Noah E. Berlow, Charles Keller, Guangheng Li

**Affiliations:** Children’s Cancer Therapy Development Institute, Beaverton, OR 97005 USA; Genetics Branch, Oncogenomics Section, Center for Cancer Research, National Cancer Institute, Gaithersburg, MD, USA; Greehey Children’s Cancer Research Institute, University of Texas Health Science Center, San Antonio, TX 78229 USA

## Abstract

Alveolar and embryonal rhabdomyosarcoma (RMS) are soft-tissue cancers that affect children, adolescents, and young adults. Sometimes referred to as muscle cancer, RMS is a cancer of muscle and non-muscle origin that phenocopies incompletely differentiated myoblasts or activated satellite (muscle stem) cells. Interestingly, embryonal RMS (ERMS) has been observed to undergo terminal myogenic differentiation in response to stress induced by chemotherapy and radiation therapy^4, 9, 24^. Given the propensity of rhabdomyosarcoma to differentiation, in this report we explore the use of differentiation therapy combining MEK inhibitor (MEKi) cobimetinib and chemotherapy as a strategy to halt RMS growth. We evaluated a representative panel of RMS cell lines with cobimetinib and chemotherapy in two dosing schedules that mimic clinical use followed by cell growth evaluation and high content analysis (differentiation) assays. We uncovered that cobimetinib does not have significant additive or synergistic effects on cell differentiation or cell growth with chemotherapy in RMS and can have unanticipated antagonistic effects; specifically, pre-exposure of cobimetinib to cells can decrease the effectiveness of chemotherapy-mediated cell growth inhibition *in vitro*. Although differentiation-therapy is still a potential viable strategy in RMS, our data do not support MEKi/chemotherapy co-treatment in this context.

## Introduction

The long-term outcome for most children with progressive/metastatic rhabdomyosarcoma (RMS) has remained dismal for more than 4 decades^21^. While temsirolimus offers a short-term survival benefit^16^, new approaches for improved long term survival benefit are needed.

Studies by Shern, Khan *et al*.^22^ have created interest in the RAS-MEK-ERK pathway for RMS. From 147 rhabdomyosarcoma samples investigated by whole genome sequencing or whole exome sequencing, 11 cases (11.7%) harbored *NRAS* mutations, 6 cases (6.4%) harbored *KRAS* mutations, and 4 cases (4.3%) harbored *HRAS* mutations. Thus, 22% of cases for fusion negative tumors (ERMS) harbor *RAS* mutations^22^; however, fusion positive alveolar RMS (ARMS) in general do not.

RAS is upstream of MEK1/2 and in turn ERK1/2^8^. Cobimetinib is a representative MEK inhibitor. Cobimetinib has been approved by the FDA for treatment of unresectable or metastaic melanoma with a BRAF mutation, in combination with vemurafenib. Cobimetininb is a MEK1-selective inhibitor that inhibits MEK1 and MEK2 with IC_50_s of 0.95 and 199 nM^31^. From clinical trial results, the recommended cobimetinib dose of 60 mg resulted in average exposures of 514 nM (C_max_)^31^.

The literature does not yet systematically address the use of MEK inhibitors in RMS, but published studies are informative for the MEK1 signaling pathway as a target. In the ERMS cell line RD (harboring an *NRAS* Q61H mutation), the MEK inhibitor U0126 induced G_1_/S cell cycle arrest and reduced cell growth 8-fold^14^. Synergy with irradiation is also observed^15^. For the same cell line, the MEK inhibitor AZD6244 slowed tumor growth with an IC_50_ of ~ 100 nM (by comparison, for the ARMS cell line Rh30, the IC_50_ for AZD6244 was > 10 uM^13^). Soberingly, AZD6244 had no affect on tumor growth as a single agent in RD xenografts despite phospho-ERK reduction in peripheral mononuclear cells. Combining MEK and AKT inhibitors synergistically further slowed (but did not stop) tumor growth^19^. These results point to an interest in MEK inhibitors for RMS, but likely in the context of better drug combinations rather than as a single agent.

We address a unique therapeutic application of MEK inhibitors as agents that halt tumor growth by the induction of terminal myogenic differentiation. Clinical evidence suggests that inducing terminal differentiation of tumor cells is a feasible goal since myogenic differentiation under stress is an intrinsic property of RMS after treatment with chemotherapy and radiation^4, 9, 24^. This effect is more prominent in ERMS than ARMS. Yohe *et al*. provide a possible explanation in the observation that the MEK inhibitor trametinib de-represses the *MYOG* promoter^33^. Albeit, these experiments were performed at 3-times the maximum achievable clinical concentration (C_max_ 36 nM, C_sss_ 21 nM^12^) and 50-100 fold higher than the K_d_ values for MEK1/2 (0.92 and 1.8 nM, respectively^32^). Nevertheless, fully harnessing embryonal and alveolar RMS differentiation potential clinically is the purpose of this study.

We have evaluated two differentiation-therapy strategies in this report. The first strategy hypothesizes that chemotherapy-induced myogenic differentiation in RMS is a 2-step process: (i) debulking the tumor cell mass with stress-inducing chemotherapy, then (ii) MEK inhibition-induced reprogramming of tumor-propagating cells (TPCs) into terminally-differentiation myogenic cells. The second strategy hypothesizes that preexposure of tumor cells to MEKi prior to chemotherapy treatment will sensitize RMS cells to chemotherapy-induced differentiation.

In our studies, we found that RMS cell lines exhibit a broad range of sensitivity to MEK inhibitors and chemotherapy agents. We also found that MEKi and chemotherapy agents can behave antagonistically in RMS; that is, treatment with a MEKi can decrease chemotherapy sensitivity. Finally, we found that MEKi and chemotherapy treatment induces myoblast differentiation but that this treatment strategy is insufficiently robust to support clinical development in RMS.

## Results

### MEK1/2 pathway components are selectively over-expressed at the RNA level

MEK1 is highly expressed in ERMS and in some ARMS biopsies. The MAPK signaling pathway components are highly expressed in selected ARMS cases as well (Figure 1). As a secondary check to simple gene expression data, pathway signatures have been assessed using S-score method described previously^1, 20^. We have already shown that 21% of ERMS have a strong “RAS-on” signature^20^ and upon examination 23% of fusion-positive ARMS have this same RAS-on signature (Figure 2A and 2B). Interestingly, MEK1 or MEK2 expression were each associated with improved survival following chemotherapy in the Intergroup Rhabdomyosarcoma Study-IV (Suppl. Figure 1).

**Fig 1.**
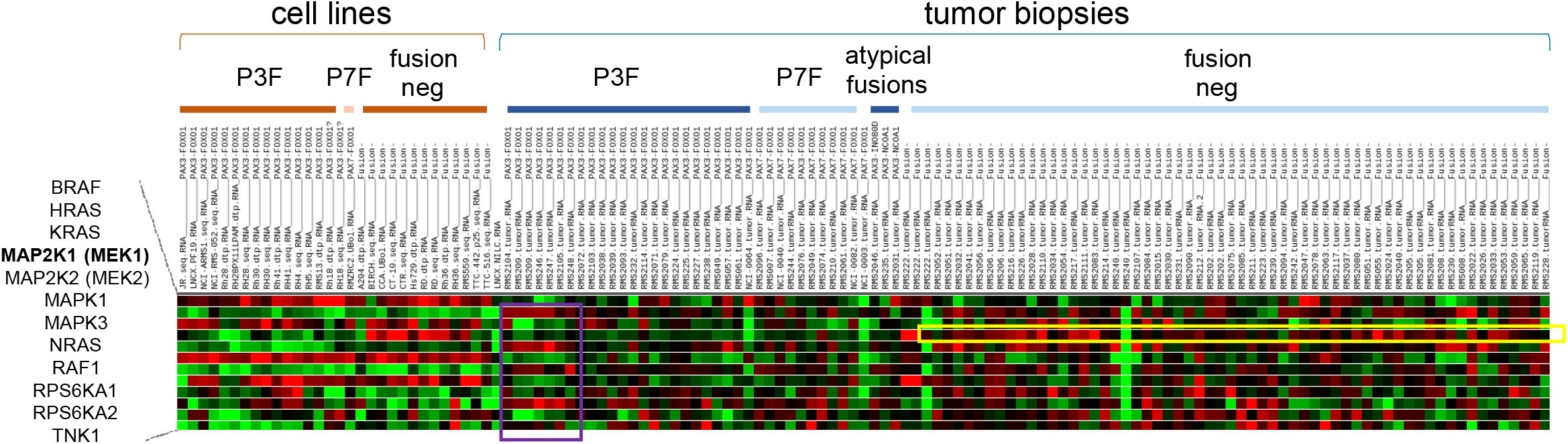
RAS signaling axis component expression in RMS (RNAseq). Yellow, MAP2K1 (MEK1) expression is most elevated across fusion negative (embryonal) RMS. Purple, select PAX3:FOXO1 ARMS tumors express elevated downstream components.

**Fig 2.**
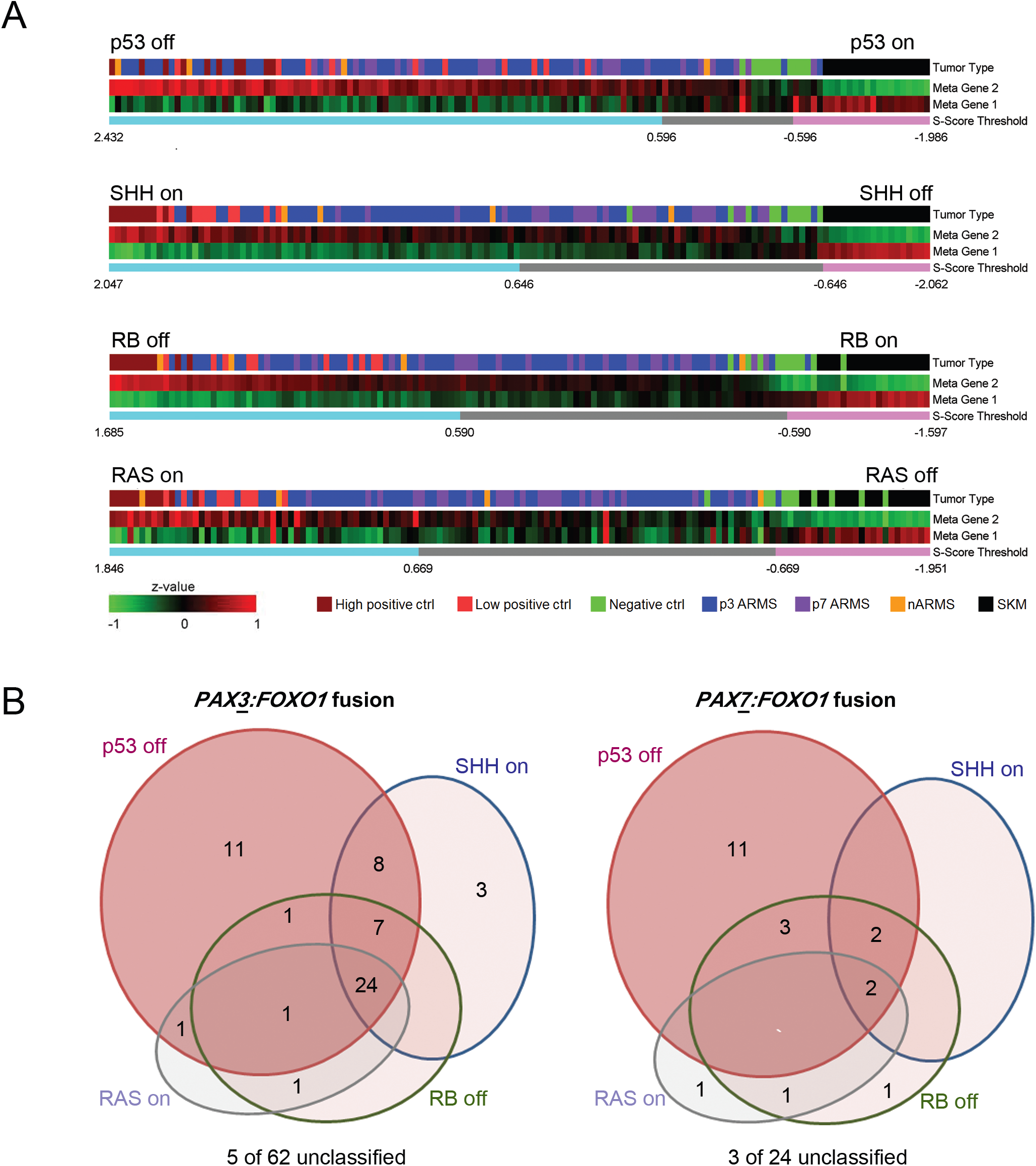
RAS is a modifier in most human RMS. (A) Heatmaps for the p53 pathway, SHH pathway, RB pathway and RAS pathway are each represented as two metagenes as previously described^20^. In total, 53 of 62 *PAX3:FOXO1*+ tumors exhibited a gene expression signature consistent with the “p53-off” state for which S-score was greater than 0.596. Eighteen of 24 Pax7:Foxo1 tumors also had a p53-off state. For SHH, 42 of 62 *PAX3:FOXO1*+ tumors and 4 of 24 *PAX7:FOXO1*+ tumors exhibited a SHH- on overdrive state for which S-score was greater than 0.646. For Retinoblastoma protein, 24 of 62 *PAX3:FOXO1*+ tumors and 9 of 24 *PAX7:FOXO1*+ tumors exhibited a RB-off state for which S-score was greater than 0.590. For RAS, 35 of 62 *PAX3:FOXO1*+ tumors and 4 of 24 *PAX7:FOXO1*+ tumors exhibited a RAS-on overdrive state for which S-score was greater than 0.669. (B) Venn diagram of signatures, revealing that p53 is the most common pathological signature overall or to occur independently of other signatures.

### MEK1/2 pathway components are selectively over-expressed at the protein level

We performed immunohistochemistry of the MEK targets ERK1/2, scoring each sample by intensity and fraction (%) cells stained (Figure 3). In brief, ERK1/2 phosphorylation is found in 36% of ERMS and 14% of fusion-driven ARMS, in both nuclear and cytoplasmic patterns, and in both tumor cells and microenvironment cells (vascular endothelium/stroma) (Figures 3A and 3B). This result is consistent with our earlier finding that both ERMS and ARMS are associated with a RAS-on signature. Immunoblotting studies were also performed to discern tumor cell vs microenvironment phosphorylation status of this MEK1 target. We also evaluated fresh RMS biopsies for phosphorylated-ERK1/2 and similar to IHC, biopsies showed variable phospho-ERK expression (Figure 3C).

**Fig 3.**
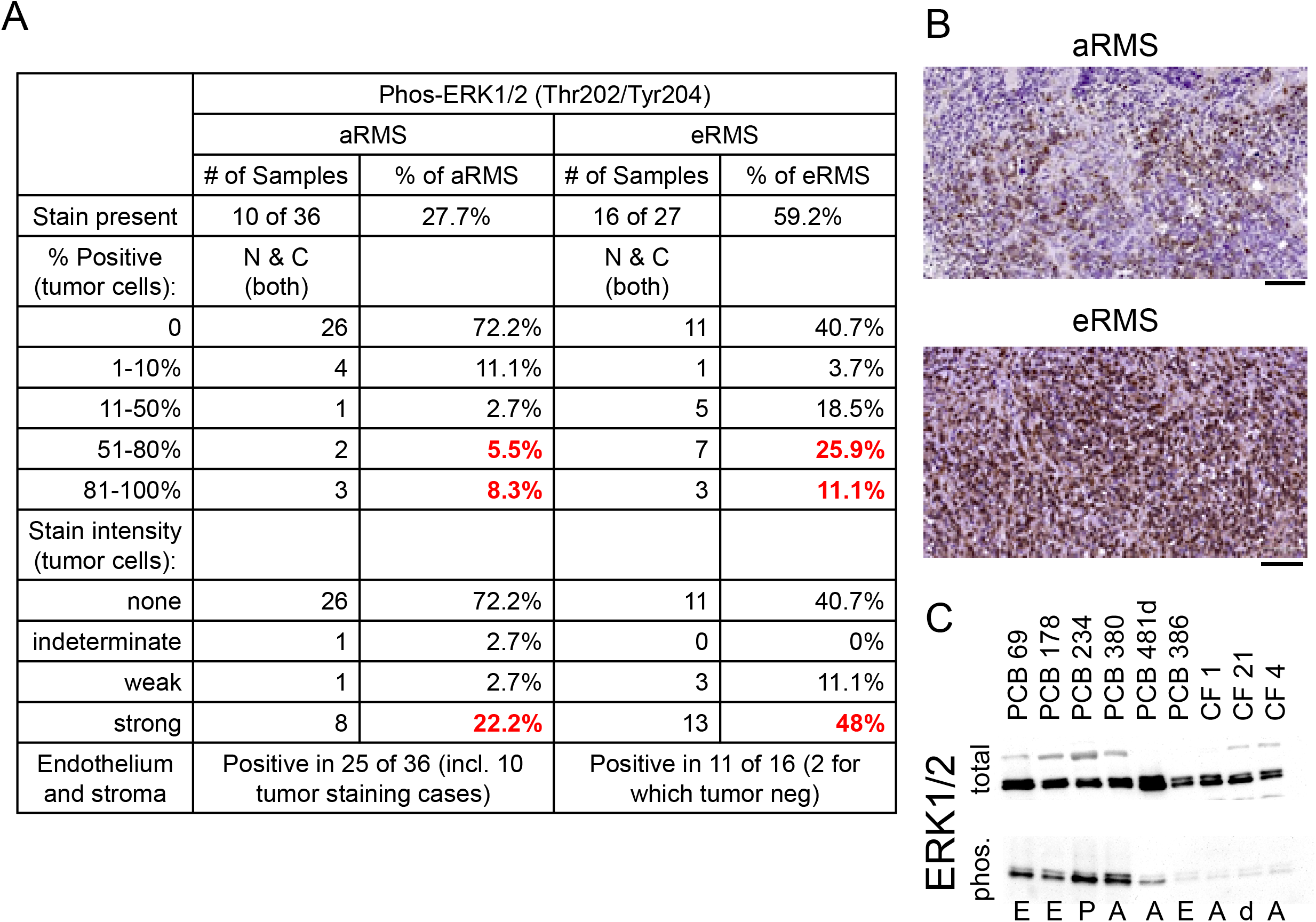
Human and GEMM TMAs for pERK (Thr202/Tyr204). **a** Quantification of tumor samples. **b** representative photomicrographs. scale bar, 100 μm. **c** Confirmatory western blot analysis of tumor biopsies. E, ERMS; A, ARMS; P, pleomorphic RMS; d, desmoplastic round cell tumor.

### RAS signaling in RMS may generally be a modifier of disease rather than a driver

For the “RAS-on” signature in ERMS^20^, and in ARMS presented above, we posited that if RAS signaling were a driver event, then a RAS-on signature would exist exclusive of other RMS pathway signatures. Using our previously described metagene analysis and S-score method^20^, we analyzed a global gene expression dataset of 62 *PAX3:FOXO1*+ and 24 *PAX7:FOXO1*+ human alveolar rhabdomyosarcoma tumors and categorized them as containing the expression signatures of rhabdomyosarcoma-associated pathological determinants: tumor suppressor protein p53 (*TP53*), sonic hedgehog (*SHH*), retinoblastoma protein (*RB1*) or *RAS* (Figure 2B). We found that 85% of *PAX3:FOXO1*+ tumors exhibited a gene expression signature consistent with the p53-off state. Similarly, 75% of *PAX7:FOXO1* tumors also had a p53-off state. For SHH, a surprising 68% of PAX3:FOXO1+ but only 17% of *PAX7:FOXO1*+ tumors exhibited a SHH-on overdrive state. For Retinoblastoma protein signature, 39% of *PAX3:FOXO1*+ and 38% of *PAX7:FOXO1*+ tumors exhibited a RB-off state. For RAS signature, 56% of *PAX3:FOXO1*+ and 17% of *PAX7:FOXO1*+ tumors exhibited a RAS-on overdrive state. This last result is consistent with the observation that *KRAS* mutation can occur in ARMS^23^ (specifically, *PAX7:FOXO1*+ ARMS), although uncommonly as a driver mutation. Thus, the p53-off state was the most common signature abnormality found either alone or in combination, while the other signatures (SHH activation, RAS activation, RB inactivation) were rarely found without a combined p53-off signature (Figure 2). This results for a RAS activation signature as a modifier of disease, but not as a driver, is similar to what we had reported for ERMS ^20^.

### Response of MEKi-alone and combination MEKi plus chemotherapy treatment of RMS cell lines in a proliferation assay

RMS cell lines are sensitive to growth arrest by MEK inhibitor cobimetinib treatment (Figure 4). However, the RMS cell lines are sensitive at a drug concentration above what can be achieved clinically (cobimetinib steady state C_max_=514nM) even for NRAS-mutation bearing RD cells (IC_50_=706nM). As a control for the assay melanoma line A2058 (BRAF V600E) was also analyzed. The IC_50_ value for A2058 (136 nM) is well below cobimetinib’s clinical C_max_ (514nM). To improve the efficacy of cobimetinib we have explored the combination with chemotherapy. For this, we tested two drug-dosing schedules, (i) 24-hour MEKi exposure followed by addition of chemotherapy for an additional 48-hour, and (ii) 48-hour chemotherapy exposure followed by addition of MEKi for an additional 24-hour (Figure 5 for vincristine; Suppl. Figure 2 for vinorelbine and mafosfamide). Dosing schedule (ii) showed little MEKi-dependent effect. Possibly, the terminal 24-hour MEKi exposure was not of sufficiently length to reveal an effect. Dosing schedule (i) did reveal a MEKi effect. Using dosing schedule (i), RMS cell lines demonstrate three types of drug response: weak response to MEKi and chemotherapy, response to chemotherapy without MEKi-induced sensitization, and MEKi-induced antagonism of chemotherapy. We note that these three types of response were independent of RMS pathology (ERMS or ARMS), and independent of chemotherapy tested. We attempted to determine a molecular marker that would correlate with the three types of response. For this, we surveyed representative cell lines for their phospho-ERK1/2 status. All cell lines evaluated have similar levels of phospho-ERK1/2 prior to drug treatment, except for Rh30, which was lower. (Figure 5C).

**Fig 4.**
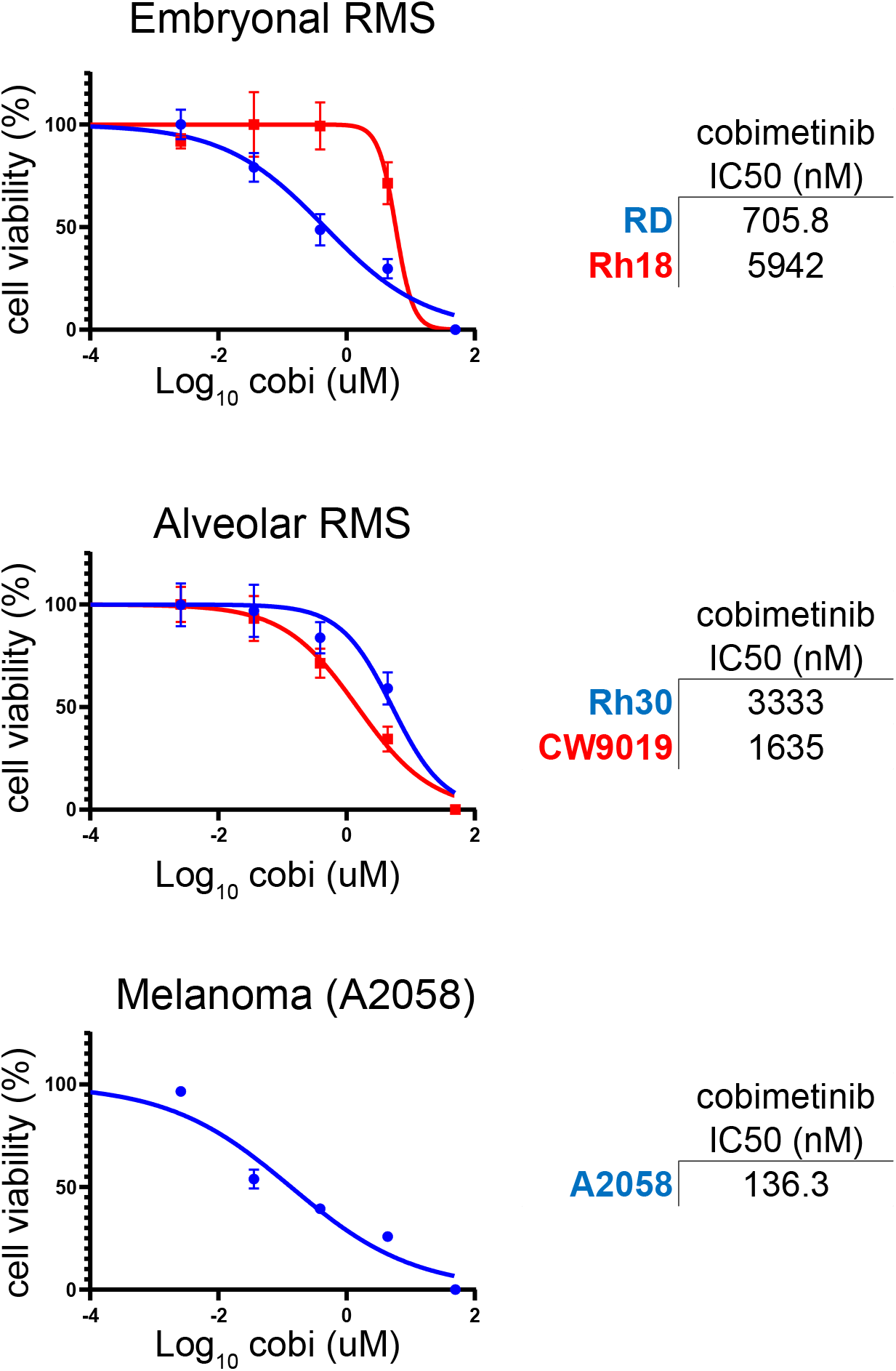
MEK inhibitor cobimetinib IC_50_ determinations for representative embryonal-, and alveolar-RMS cell lines. IC_50_ values are average of three independent experiments. Melanoma line A2058 (BRAF V600E) was included as an assay control.

**Fig 5.**
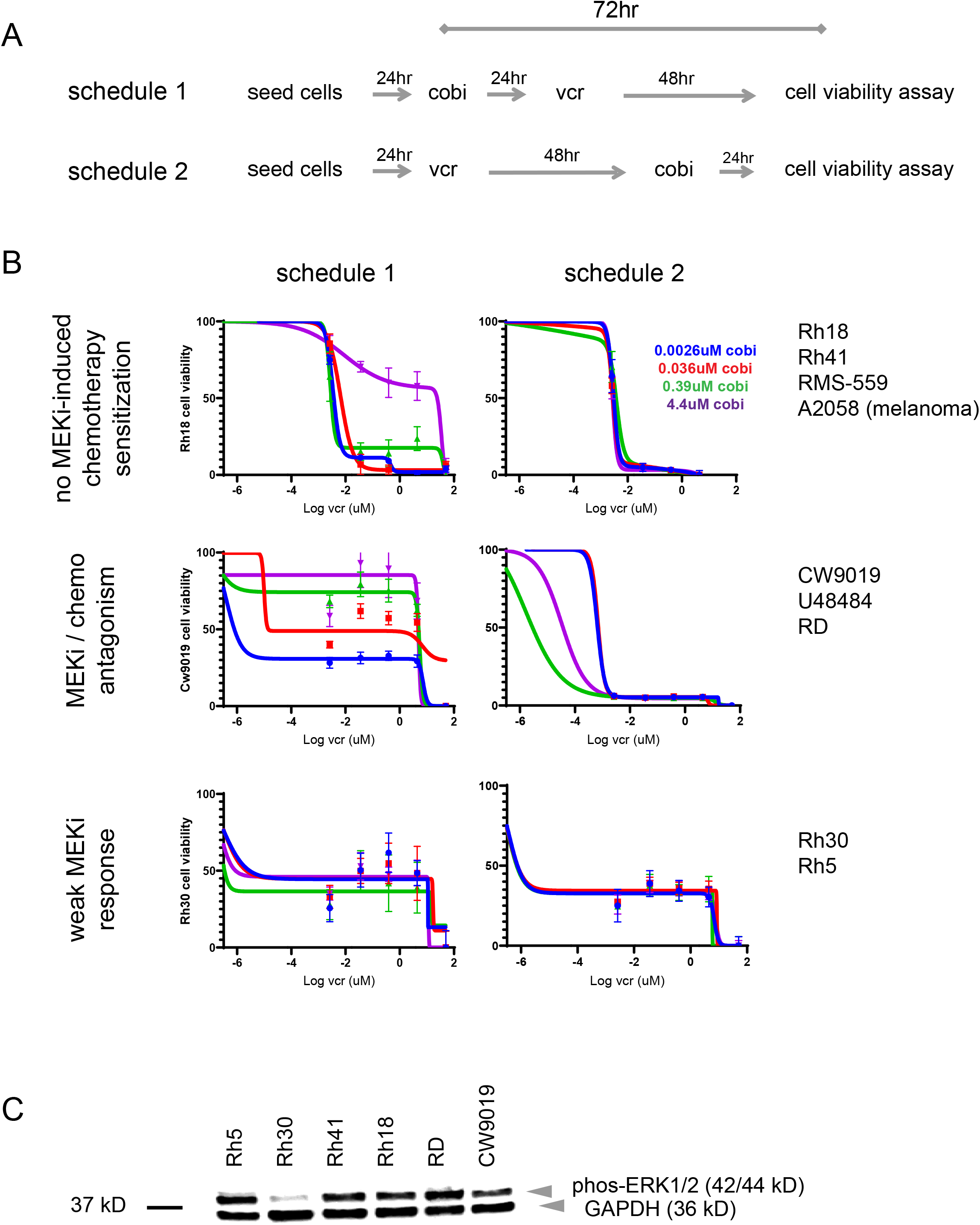
Combinations of MEKi plus chemotherapy in cellular proliferation studies. **a** Schematic of two dosing schedules. **b** Cell viability curves that illustrate three types of observed responses. **c** Baseline pretreatment phospho-ERK1/2 western blotting analysis of cell lines representing the three types of responses. hr, hours; cobi, cobimetinib; vcr, vincristine; kD, kDal.

### Response of MEKi-alone and combination MEKi plus chemotherapy treatment of RMS cell lines in a differentiation assay

We further evaluated MEKi (cobimetinib) and chemotherapy (vincristine) treatment in a myogenic differentiation assay (Figure 6 and Supple. Figure 3). For this, human RMS cell lines RD (ERMS) and Rh30 (ARMS) and mouse myoblast line C2C12 were cultured at high cell-density and low-serum (2% horse serum) for three days prior to exposure of MEKi. Following MEKi exposure, the cultures were then exposed to vincristine chemotherapy and assayed for markers of proliferation (Ki-67) and apoptosis (cleaved-caspase 3). We further assayed for markers of differentiation by determining the fusion-index (number of myosin heavy chain positive cells per total number of cells)), and myogenic-index (numbers of cells fused per total number of cells). Regarding cellular proliferation, we did not observe a strong cobimetinib-induced anti-proliferation effect for any of the cell lines as a single agent or any cobimetinib-induced sensitivity to chemotherapy. Regarding the apoptosis marker, the RD cell line has an elevated cleaved-caspase 3 response to vincristine; however, this signal diminishes if the cells are pre-exposed to cobimetinib. This result is again similar to MEKi/chemotherapy antagonism previously observed. For the Rh30 cell line, cobimetinib treatment as a single agent induced apoptosis and increased sensitivity to chemotherapy. This result is in contrast to what was observed in the cellular proliferation assay in which Rh30 was relatively resistant to cobimetinib and chemotherapy. This difference may reflect the differentiation state of the Rh30 culture induced by low-serum culturing. Regarding the differentiation marker (MHC), Rh30 did not exhibit the hallmarks of differentiation. In contrast, for RD cobimetinib alone led to an increase in the fusion-index. However, chemotherapy decreased the fusion index and when both drugs were administered little differentiation was seen.

**Fig 6.**
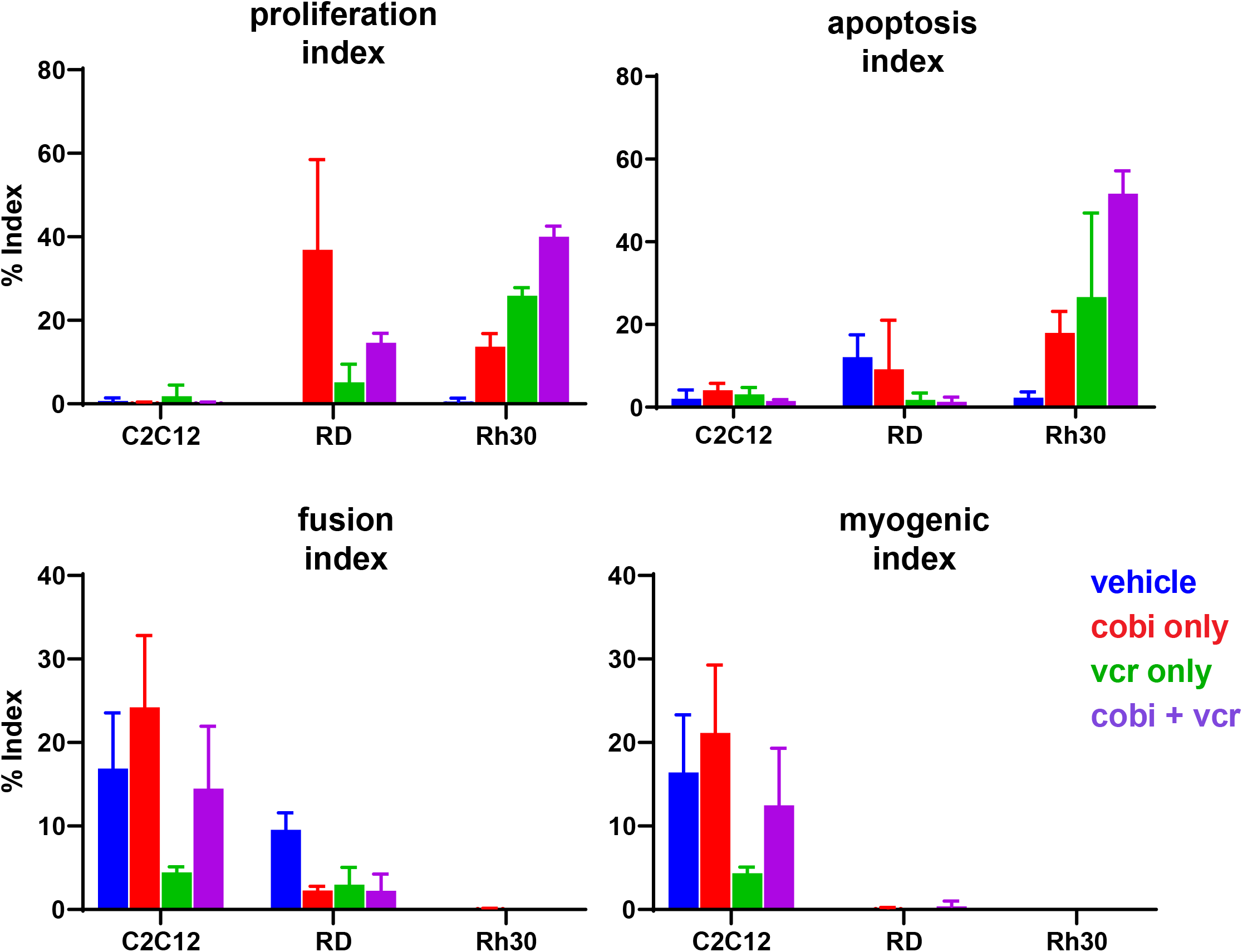
High-content analysis immunocytochemistry staining quantification. Proliferation index is the percentage of KI67 positive cells to total number of cells; apoptosis index is the percentage of cleaved caspase 3 cells to the total number of cells; fusion index is the ratio of nuclei per cell or myotube; myogenic index is the ratio of MF20(MHC)-positive cells to total cell number.

## Discussion

Myogenic differentiation as a treatment strategy for rhabdomyosarcoma is a reasonable approach^27^. Whereas RAS-ERK(MEK) signaling and chemotherapy-induced myogenic differentiation are, separately, well-recognized properties of ERMS, our data show that MEK inhibition does not enhance, and can be antagonistic to, chemotherapy-induced proliferation arrest and myogenic differentiation *in vitro*. Our data reveal that although MEK inhibition can induce many of the hallmarks of differentiation, such as myotube formation, this induction is limited in scope. MEKi treatment failed to induce robust myotube formation or robust myosin heavy chain expression despite applying non-physiological conditions such as low serum culturing to encourage differentiation.

Whether RAS is a key target in RMS is strongly debated: our gene expression, immunohistochemistry, and biochemical surveys indicate an active RAS-ERK signaling axis in up to 36-59% of ERMS and 14-28% of ARMS. However, RAS is rarely a sole mutation or sole mutational signature in either ERMS or ARMS arguing that in non-hereditary RMS, RAS may be a cooperating initiating mutation, a modifier of disease, or a co-driver (not a central driver, single point of failure). Incomplete, unsustained response to MEK inhibitors in xenograft models are consistent with this view^13^. The correlation between high MEK1/2 expression and <u>increased</u> survival in IRS-IV clinical samples is perhaps the strongest reason to question the RAS-MEK axis in RMS (Supple Fig1).

MEK inhibitors in general, and cobimetinib specifically, have been reported to work through G1 phase cell cycle arrest in several cancer types^7, 17, 26^. In contrast, chemotherapy agent vincristine is an inhibitor of microtube formation that primarily impacts chromosome separation during metaphase. Rather than working synergistically to induce differentiation as originally envisioned, our data could be interpreted as consistent with cobimetinib and vincristine mechanistically working as cell cycle inhibitors. That is, for some RMS cell lines pretreatment with cobimetinib induced G1-arrest that then reduced the effectiveness of vincristine that works later in the cell cycle. Nevertheless, neither mafosfamide (cyclophosphamide) or vinorelbine were synergistic with cobimetinib either.

Although differentiation therapy of RMS may be possible, our data indicates that combination MEKi/chemotherapy treatment in this schedule is insufficiently robust to warrant clinical development in this disease.

## Materials and methods

### Cell Lines

Cell line RD (Cat# CCL-136) and C2C12 (Cat# CRL-1772) were obtained from ATCC (Manassas, VA) and was cultured in Dulbecco Modified Eagle Media (DMEM, Thermo Cat# 11995065, Waltham, MA) supplemented with 10% FBS (Thermo Cat# 10437036) and 1% penicillin/streptomycin (Thermo Cat# 15140122) (DMEM+10:1). Rh5, Rh18, Rh30, and Rh41 cell lines were obtained from Dr P. Houghton of St. Jude Children’s Research Hospital, Memphis, TN. These lines are currently available from Childhood Cancer Repository (www.cccells.org). These lines were cultured in Roswell Park Memorial Institute media (RPMI-1640, Thermo Cat# 11875093) supplemented with 10% FBS and 1% penicillin/streptomycin (RPMI+10:1). Cell line CW9019 was obtained from the laboratory of Dr. Frederick Barr (NIH, Bethesda, MD) and this line was cultured in DMEM+10:1. Cell line RMS559 was obtained from the laboratory of Dr. J. Fletcher (Brigham and Women’s Hospital, Boston, MA) and was cultured in DMEM+10:1. The murine cell line U48484 was derived from a *Myf6^ICNm/WT^ Pax3^P3Fm/P3Fm^ Trp53^F2-10/F2-10^* transgenic mouse, previously described^2^. U48484 was cultured in DMEM+10:1. The melanoma cell line A2058 was obtained from ATCC (ATCC CRL-11147) and was cultured in DMEM+10:1.

### Western blot analysis

For western blot analysis, protein lysates were prepared using a standard RIPA buffer (Thermo Cat# 89900) and cell scrapping method. The RIPA buffer was supplemented with HALT protease and phosphatase inhibitor cocktail (Thermo Cat# 78441). Protein lysates were sampled for bicinchoninic acid assay (BCA) (Thermo Cat# 23227) to determine protein concentrations. 25ug of each protein lysate were diluted into 3xSDS gel loading buffer (New England Biolabs Cat# B7703S, Lpswich, MA) and heated to 95°C for 5 minutes. The denatured samples were loaded onto a 10% Tris-Glycine SDS-polyacrylamide gel (Bio-Rad Cat# 456-1024, Hercules, CA). Protein molecular weight marker (Bio-RAD Cat# 161-0374) was included. After gel electrophoresis, proteins were transferred to PVDF membrane (Bio-Rad Cat# 1620177) using a trans-blot method (Bio-Rad Cat# 1658030). Membranes were incubated in blocking buffer at room-temperature for 1 hour; either 5% bovine serum albumin (Jackson ImmunoResearch Cat# 001-000-162, West Grove, PA) for phosphoproteins western blots or 5% milk for all others. Blocking buffers were formulated in TBST. After blocking, membranes were incubated overnight at 4°C, rocking, in primary antibody diluted in fresh blocking buffer. The following primary antibodies were used: p44/42 MAPK (Erk1/2) (Cell Signaling Cat# 9102, Danvers, MA), Phospho-p44/42 MAPK (Erk1/2) (Thr202/Tyr204) (Cell Signaling Cat# 9101), ß-Actin (13E5) (Cell Signaling Cat# 4970). After appropriate washing, HRP-conjugated secondary antibody (Vector Laboratories Cat# PI-1000, Burlington, CA) incubation was performed at room temperature for 2 hours. Following antibody incubations and washes, the blot was treated with chemiluminescent substrates (Bio-Rad Cat# 1705061) and visualized on a FluorChem™ Q imaging system (ProteinSimple).

### Sequence analysis

DNA and RNA sequencing methodologies were previously published (Nat. Med. 2015 June; 21 (6): 555-559). Data were obtained from the OncoGenomic DB maintained by the laboratory of Javed Khan (National Cancer Institute).

### Microarray analysis for S-Scores and Principal Component Analysis

Human microarray datasets were previously published^6, 10, 30^. Patient demographics are presented in Supplemental Tables 1a and 1b. Metagene and S-score analysis were conducted as previously described^20^. For mouse tumors, gene expression analysis was performed using Illumina’s Mouse Ref 8 Beadchip v1 (Illumina Inc, San Diego, CA). These datasets have been deposited in GEO as accession entry GSE22520 and described in Supplemental Table 1c. Rank invariant normalization was performed on the log_2_-transformed expression value. Afterwards, we applied Principal Component Analysis (PCA) to all mouse tissue samples with the 19,070 probes (selected with the criterion of average log_2_-intensity > 5.5 and standard deviation > 0.1) and then plotted in 3D space for visualization (Figure 4b). Similar microarray data analysis and the PCA methods were previously described ^20^. All bioinformatics tasks were performed with MATLAB/Bioinformatics Toolbox (MathWorks Inc, Natick, MA) unless otherwise noted.

We downloaded the public domain datasets for fusion negative rhabdomyosarcoma reported by Davicioni *et al*^5^ from (https://array.nci.nih.gov/caarray/project/details.action?project.experiment.publicIdentifier=trich-00099) as well as ARMS rhabdomyosarcoma datasets from previously published reports ^10, 30^. These fusion negative RMS and ARMS datasets were designated as the test samples, whereas normal skeletal muscles (SKM) samples reported by Bakay *et al*.^3^ were used as the control group. We also downloaded signature specific datasets as described in the paragraphs below. All the studies had been performed on Affymetrix U133A array platform (Affymetrix, Santa Clara, CA). Sample IDs used are given in Supplemental Tables 1 and 2. Control samples of each examination are the embryonal rhabdomyosarcoma (ERMS) samples derived from Rubin *et al*.^20^. We have selected 10 high positive, 10 low positive and 10 negative control samples for each examination, whereas gene-wise t-tests were performed between the 20 positive controls and normal SKM in order to apply to S-score. S-score is a subtype scoring method to quantify each sample’s consistency. By using the S-score, we can unambiguously identify sample status and the amplitude of pathways or biological processes that gene signatures have depicted. The detailed S-score method has been previously described^20^.

**Table 1.**
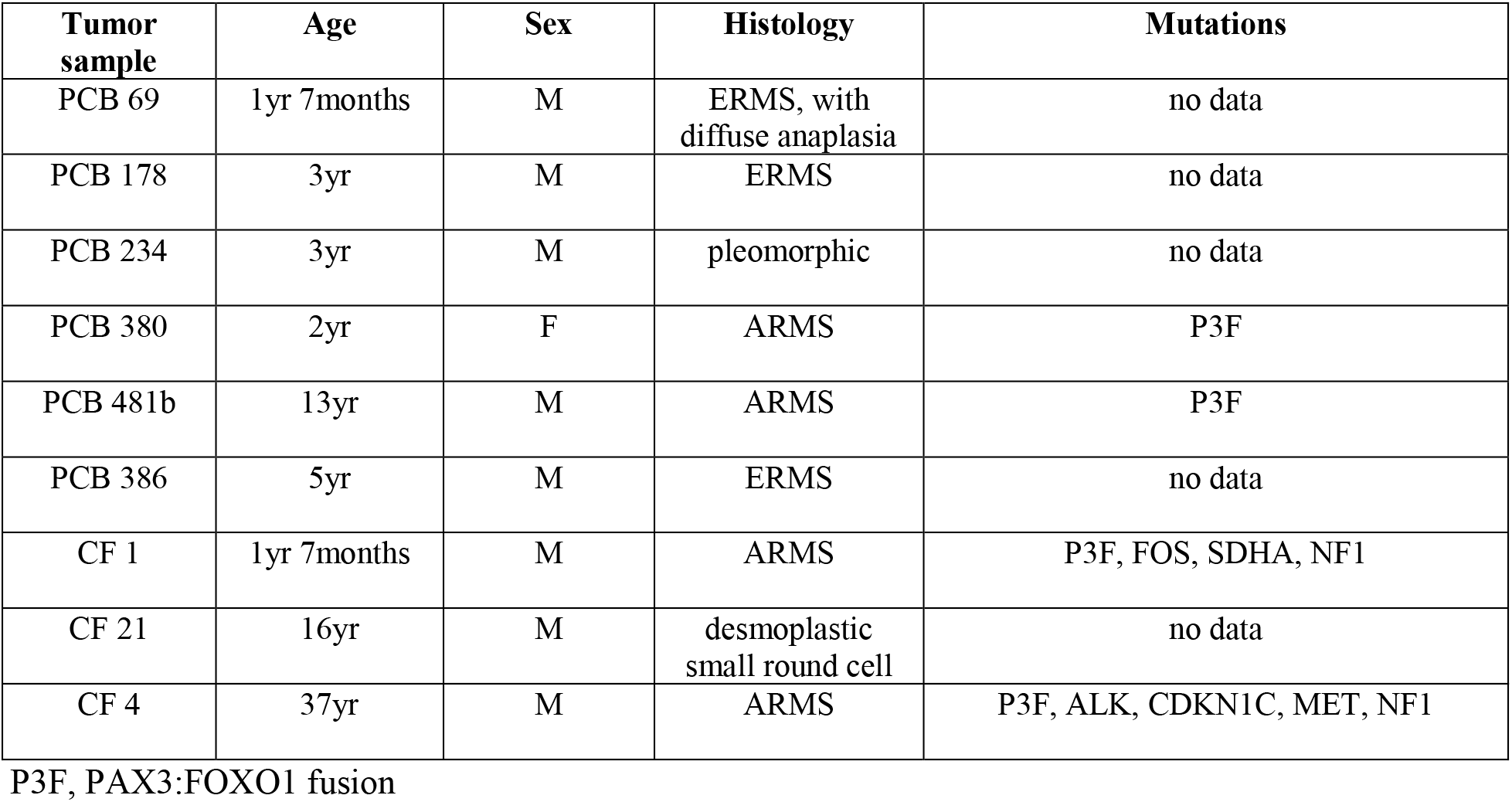
Tumor samples evaluated for ERK protein status

| **Tumor sample** | **Age** | **Sex** | **Histology** | **Mutations** |
| --- | --- | --- | --- | --- |
| PCB 69 | 1yr 7months | M | ERMS, with diffuse anaplasia | no data |
| PCB 178 | 3yr | M | ERMS | no data |
| PCB 234 | 3yr | M | pleomorphic | no data |
| PCB 380 | 2yr | F | ARMS | P3F |
| PCB 481b | 13yr | M | ARMS | P3F |
| PCB 386 | 5yr | M | ERMS | no data |
| CF 1 | 1yr 7months | M | ARMS | P3F, FOS, SDHA, NF1 |
| CF 21 | 16yr | M | desmoplastic small round cell | no data |
| CF 4 | 37yr | M | ARMS | P3F, ALK, CDKN1C, MET, NF1 |

P3F, PAX3:FOXO1 fusion

**Table 2.**
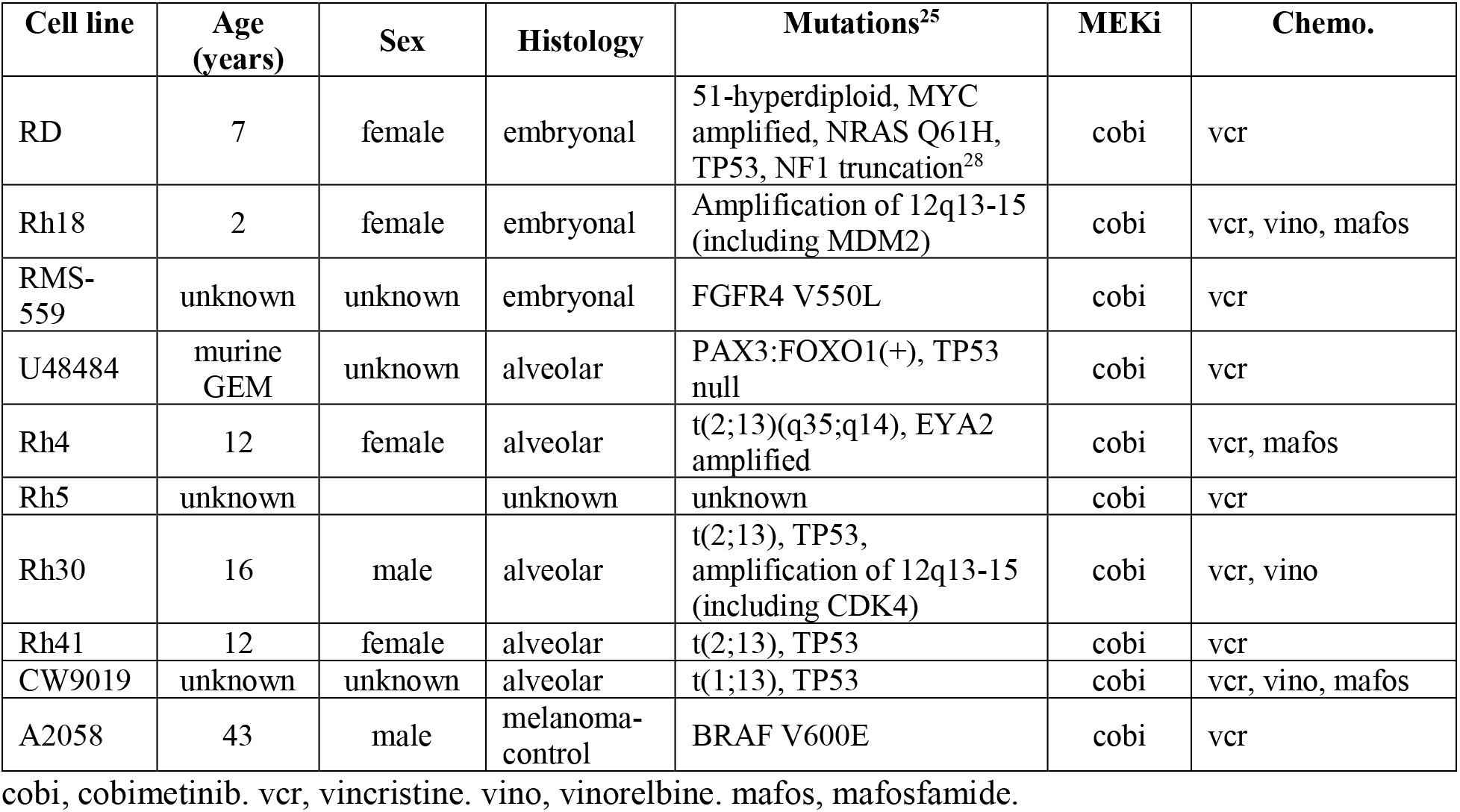
Cell lines evaluated in combination drug-screens

We used previously established gene lists^20^ to examine whether human fusion negative RMS and ARMS tumors had evidence of p53 loss of function, SHH gain of function, RB loss of function, or RAS activation. Gene signatures of p53 loss of function were derived from gene expression dataset in breast cancer^18^. We also downloaded datasets for medulloblastoma samples known to exhibit a SHH gain of function signature^29^. We took homolog genes from *Rb1* wildtype and homozygous *Rb1* deleted fusion-negative mouse sarcomas to be RB loss of function gene signature^20^. For Ras activation, we used gene lists for the activated RAS signature of zebrafish ERMS^11^. The details of obtaining gene signature of each case have been previously described^20^.

### Tissue microarray analysis

Tissue microarray analysis was performed on 2 mm tissue punches that were formalin-fixed and paraffin-embedded. The specimens were histological evaluated for rhabdomyosarcoma sub-type. Specimens were probed with phospho-p44/42 MAPK (ERK1/2) (Thr202/Tyr204) (20G11) antibody (Cell Signaling Cat# 4376).

### Cellular proliferation assay

MEK inhibitor (MEKi) and chemotherapy reagents were evaluated in a cellular proliferation assay. Both drugs were formulated to 10mM in DMSO. Rhabdomyosarcoma cell lines were seeded in 384-well plates at 2,500 cells per well, in 50 μL of media, using a Thermo Scientific™ Matrix Wellmate™ dispenser using a 10 μL tubing cassette. Two drug dosing schedules were evaluated in this study. In schedule 1, the day after cell seeding, cells were treated with MEKi at six concentrations of a log-based dilution series (0, 2.6 nM, 36 nM, 0.39 μM, 4.4 μM, and 50 μM). Drug was administered with a Tecan (Mannedorf, Switzerland) D300e drug printer using a T8+ Dispensehead cassette. Following 24-hour MEKi treatment, chemotherapy drug was administered using the same protocol to generate a complete matrix of MEKi/Chemotherapy therapy. Cultures were then incubated for an additional 72 hours using standard conditions. For drug schedule 2, the day after cell seeding, the plates were treated with chemotherapy. After 48 hours of culturing the plates were then treated with MEKi. After 72 hours of complete drug treatment, cultures were treated with an equal volume of CellTiter-Glo© reagent (Promega Cat# 2020-01-29, Madison, WI). Plates were incubated for 10 minutes at room temperature, rocking while light-shielded. After which, luminescence was measured in a Biotek (Shoreline, WA) plate reader. The data was then analyzed using Prism (San Diego, CA) Graphpad© software.

### Myotube differentiation assay

Cobimetinib (cobi) and vincristine (vcr) co-treatment was evaluated in a differentiation assay. Cell lines RD (Embryonic RMS) and Rh30 (Alveolar RMS) were seeded in 96-well plate at 5,000 cells per well. After 24 hours of incubation the media was changed to low serum (2% horse serum, Thermo Cat# 16050114) media to induce myoblast differentiation. The cultures were incubated for 72 hours. After which, cobimetinib was administered for two days at 500 nM. This was accomplished with daily media changes. After this treatment the cultures were exposed to 150 nM vincristine for three days. The cultures were then formalin fixed and analyzed by ICC for markers of differentiation (Myosin 4), proliferation (Ki-67), apoptosis (cleaved caspase 3), as well as Hoechst 33842 (Thermo Cat# 41399) nuclear staining. The Myosin 4 was visualized with antibody MF20 Alex Fluor 488 (Thermo Cat# 53-6503-82), Ki-67 was visualized with monoclonal antibody (Thermo Cat #14-5698-80), cleaved caspase 3 was visualized with antibody (Promega Cat # G7481)) labeled with Qdot 655 Streptavidin conjugate (Thermo Cat# Q10121MP). Samples were then analyzed using Thermo Scientific™ Cellnsight™ High Content Screening platform (CX5110).

## Acknowledgements

This work was supported in part by a sponsored research agreement with Roche-Genentech, as well as a research grant from the Aiden’s Army Fund of St. Baldrick’s Foundation. We thank Alexandria Harrold (CC-TDI) Stephen Simko and Ana Rodriguez of Genentech’s Innovative Pediatric Oncology Drug Development (iPODD) and Yidong Chen (UTHSCSA) for their technical assistance.

## Compliance with ethical standards

All human samples were de-identified from the cc-TDI CureFast registry. The CureFast registry obtains IRB-approved informed consent for each enrollee.

## Additional Information

### Data Deposition and Access

Not applicable.

### Author Contributions

Designed studies: KAC, SS, AR, GL, CK

Performed experiments: KAC, MMC, CAR, HIHC, MNS

Analyzed data: KAC, MMC, CAR, JFS, JK, HIHC, CK, NEB

Wrote manuscript: KAC, GL, CK

## Abbreviations

ATCC: American Tissue Culture Collection
DMSO: Dimethyl sulfoxide
ERK1/2: ERK1 and ERK2
FBS: Fetal bovine serum
HRP: Horseradish peroxidase
ICC: Immunocytochemistry
IHC: Immunohistochemistry
MEK1/2: MEK1 and MEK2
PBS: Phosphate buffered saline
Phospho: Phosphorylated
PVDF: Polyvinylidene difluoride membrane
RIPA: Radio immunoprecipitation
RPM: Revolution per minute
SDS: Sodium dodecyl sulfate detergent
TBST: Tris-buffered saline with Tween 20

## Supplementary Figure Legends

**Suppl. Figure 1.**
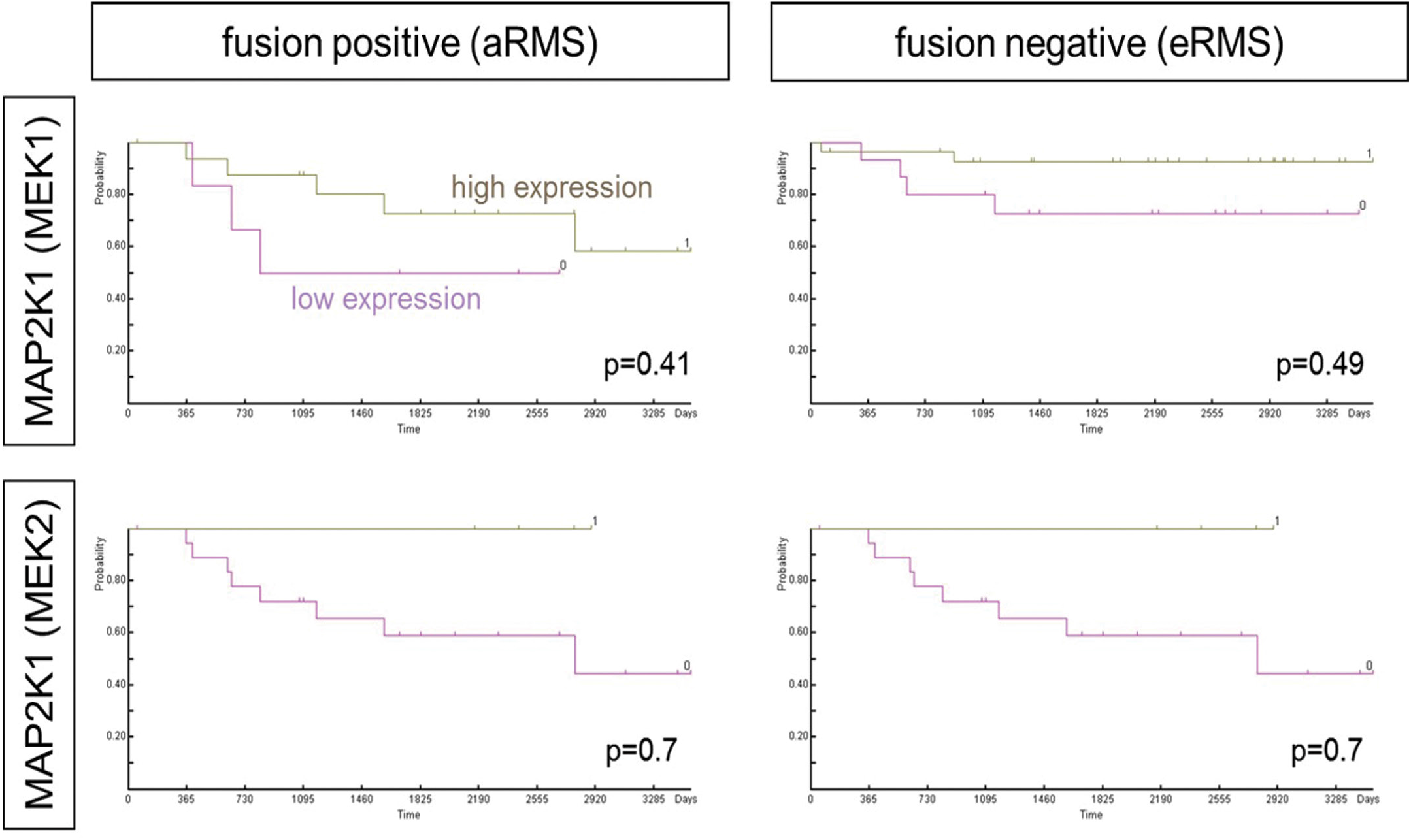
Clinical survival Vs. gene expression when adjusted for known clinical covariates (age, sex, stage, site, histology). Analysis as per [PMID 16261596].

**Suppl. Figure 2.**
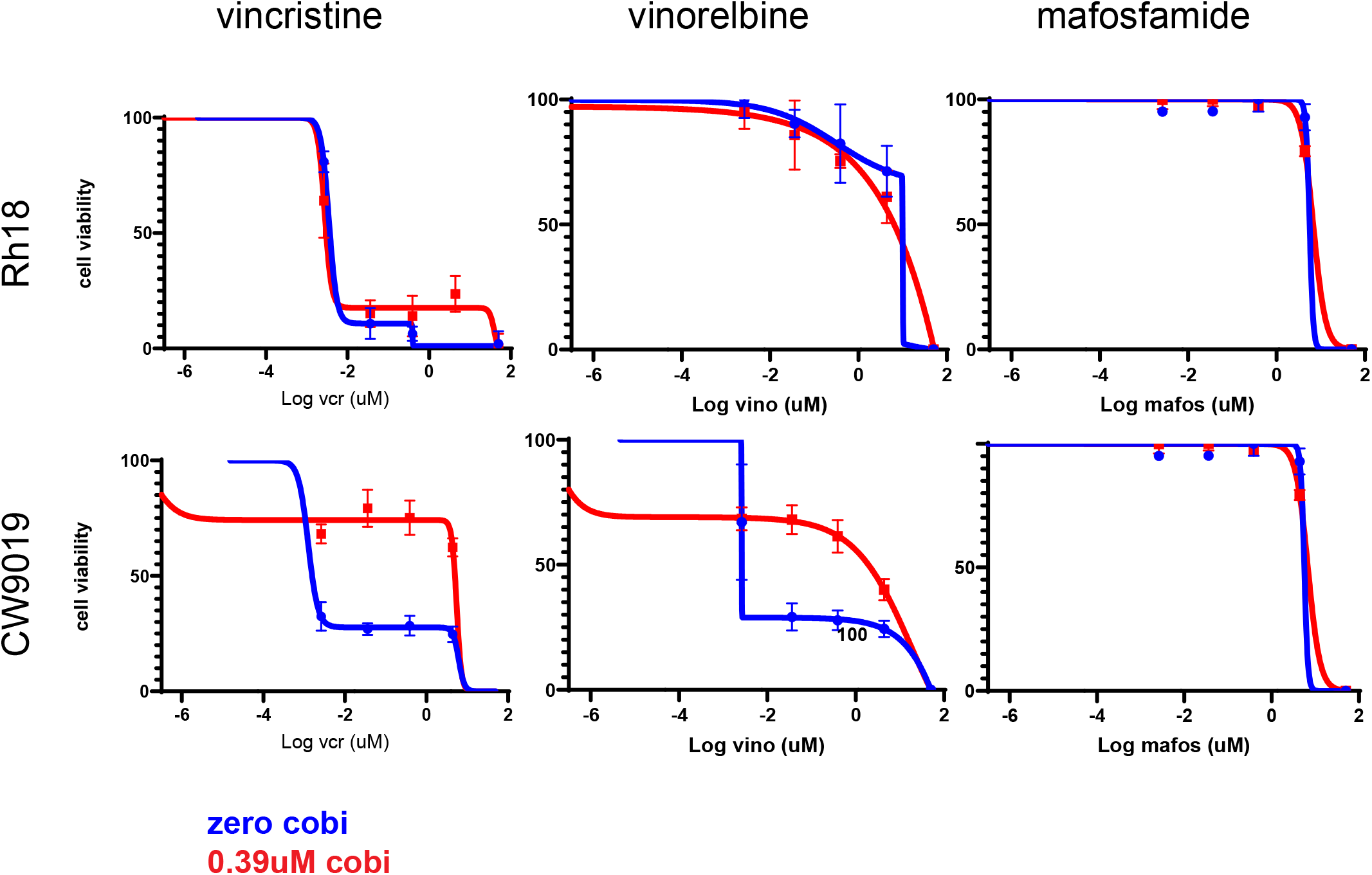
Examples of combination treatment of MEKi and chemotherapeutic agents used in relapsed rhabdomyosarcoma. Additional cell lines evaluated include Rh4 (mafosfamide) and Rh30 (vinorelbine), data not shown.

**Suppl. Figure 3.**
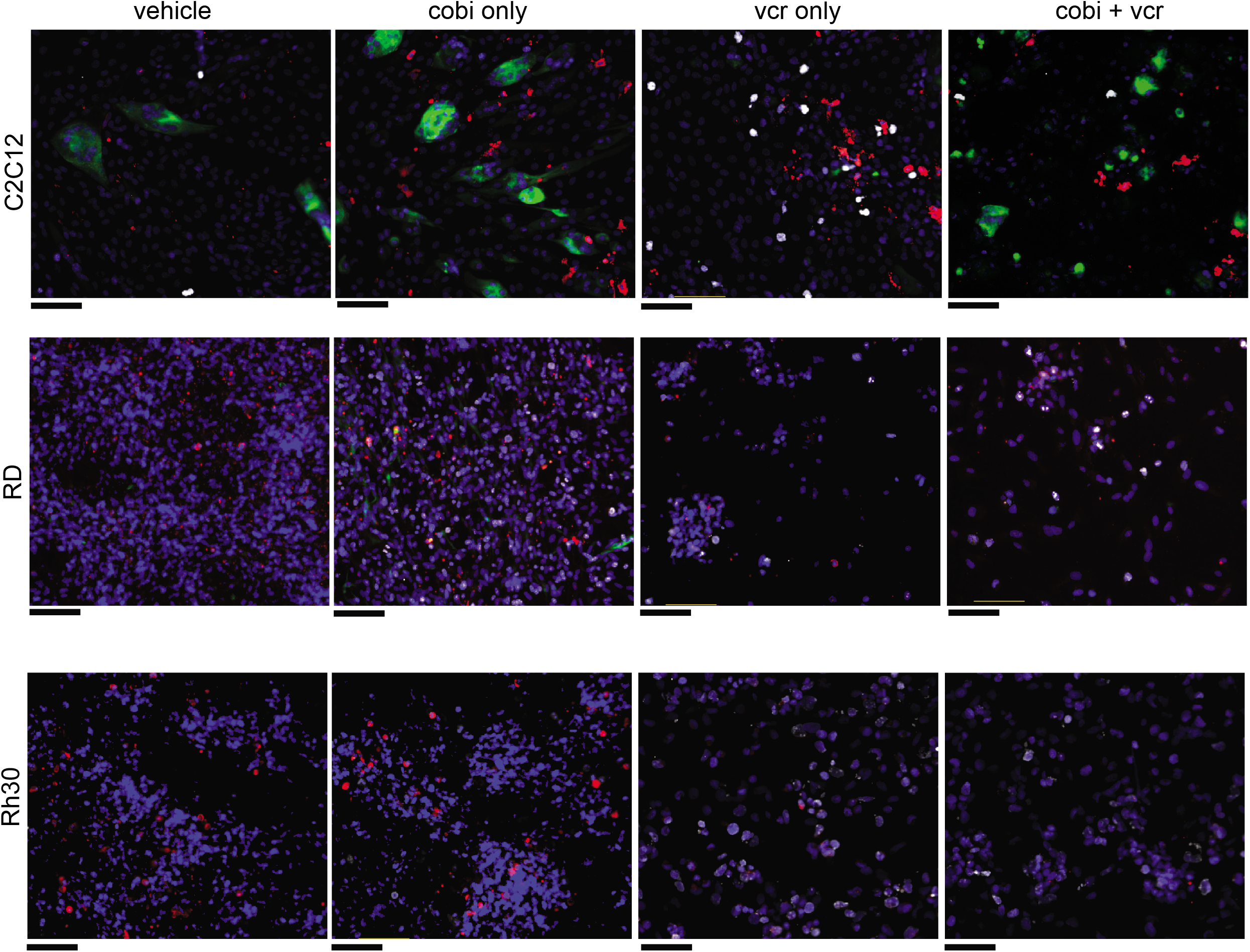
Representative images of IHC staining used to calculate graphs in Figure 6. Blue, Huechst 33842; Green, Myosin 4; Red, cleaved-caspase 3; White, Ki-67.

## References

1 Abraham J, Nunez-Alvarez Y, Hettmer S, Carrio E, Chen HI, Nishijo K et al. Lineage of origin in rhabdomyosarcoma informs pharmacological response. Genes & development 2014; 28: 1578–1591.

2 Aslam MI, Abraham J, Mansoor A, Druker BJ, Tyner JW, Keller C. PDGFRbeta reverses EphB4 signaling in alveolar rhabdomyosarcoma. Proc Natl Acad Sci U S A 2014; 111: 6383–6388.

3 Bakay M, Chen YW, Borup R, Zhao P, Nagaraju K, Hoffman EP. Sources of variability and effect of experimental approach on expression profiling data interpretation. BMC Bioinformatics 2002; 3: 4.

4 Coffin CM, Rulon J, Smith L, Bruggers C, White FV. Pathologic features of rhabdomyosarcoma before and after treatment: a clinicopathologic and immunohistochemical analysis. Modern pathology: an official journal of the United States and Canadian Academy of Pathology, Inc 1997; 10: 1175–1187.

5 Davicioni E, Finckenstein FG, Shahbazian V, Buckley JD, Triche TJ, Anderson MJ. Identification of a PAX-FKHR gene expression signature that defines molecular classes and determines the prognosis of alveolar rhabdomyosarcomas. Cancer Res 2006; 66: 6936–6946.

6 Davicioni E, Anderson MJ, Finckenstein FG, Lynch JC, Qualman SJ, Shimada H et al. Molecular classification of rhabdomyosarcoma--genotypic and phenotypic determinants of diagnosis: a report from the Children’s Oncology Group. Am J Pathol 2009; 174: 550–564.

7 Gong S, Xu D, Zhu J, Zou F, Peng R. Efficacy of the MEK Inhibitor Cobimetinib and its Potential Application to Colorectal Cancer Cells. Cellular physiology and biochemistry: international journal of experimental cellular physiology, biochemistry, and pharmacology 2018; 47: 680–693.

8 Kakadia S, Yarlagadda N, Awad R, Kundranda M, Niu J, Naraev B et al. Mechanisms of resistance to BRAF and MEK inhibitors and clinical update of US Food and Drug Administration-approved targeted therapy in advanced melanoma. OncoTargets and therapy 2018; 11: 7095–7107.

9 Keller C, Guttridge DC. Mechanisms of impaired differentiation in rhabdomyosarcoma. The FEBS journal 2013; 280: 4323–4334.

10 Lae M, Ahn EH, Mercado GE, Chuai S, Edgar M, Pawel BR et al. Global gene expression profiling of PAX-FKHR fusion-positive alveolar and PAX-FKHR fusion-negative embryonal rhabdomyosarcomas. J Pathol 2007; 212: 143–151.

11 Langenau DM, Keefe MD, Storer NY, Guyon JR, Kutok JL, Le X et al. Effects of RAS on the genesis of embryonal rhabdomyosarcoma. Genes & development 2007; 21: 1382–1395.

12 Leonowens C, Pendry C, Bauman J, Young GC, Ho M, Henriquez F et al. Concomitant oral and intravenous pharmacokinetics of trametinib, a MEK inhibitor, in subjects with solid tumours. British journal of clinical pharmacology 2014; 78: 524–532.

13 Li Z, Zhang Y, Ramanujan K, Ma Y, Kirsch DG, Glass DJ. Oncogenic NRAS, required for pathogenesis of embryonic rhabdomyosarcoma, relies upon the HMGA2-IGF2BP2 pathway. Cancer Res 2013; 73: 3041–3050.

14 Marampon F, Ciccarelli C, Zani BM. Down-regulation of c-Myc following MEK/ERK inhibition halts the expression of malignant phenotype in rhabdomyosarcoma and in non muscle-derived human tumors. Mol Cancer 2006; 5: 31.

15 Marampon F, Gravina GL, Di Rocco A, Bonfili P, Di Staso M, Fardella C et al. MEK/ERK inhibitor U0126 increases the radiosensitivity of rhabdomyosarcoma cells in vitro and in vivo by downregulating growth and DNA repair signals. Mol Cancer Ther 2011; 10: 159–168.

16 Mascarenhas L, Meyer WH, Lyden E, Rodeberg DA, Indelicato DJ, Linardic CM et al. Randomized phase II trial of bevacizumab and temsirolimus in combination with vinorelbine (V) and cyclophosphamide (C) for first relapse/disease progression of rhabdomyosarcoma (RMS): A report from the Children’s Oncology Group (COG) 2014; 32: 10003–10003.

17 Matsui TA, Murata H, Sowa Y, Sakabe T, Koto K, Horie N et al. A novel MEK1/2 inhibitor induces G1/S cell cycle arrest in human fibrosarcoma cells. Oncology reports 2010; 24: 329–333.

18 Miller LD, Smeds J, George J, Vega VB, Vergara L, Ploner A et al. An expression signature for p53 status in human breast cancer predicts mutation status, transcriptional effects, and patient survival. Proc Natl Acad Sci U S A 2005; 102: 13550–13555.

19 Renshaw J, Taylor KR, Bishop R, Valenti M, De Haven Brandon A, Gowan S et al. Dual blockade of the PI3K/AKT/mTOR (AZD8055) and RAS/MEK/ERK (AZD6244) pathways synergistically inhibits rhabdomyosarcoma cell growth in vitro and in vivo. Clin Cancer Res 2013; 19: 5940–5951.

20 Rubin BP, Nishijo K, Chen HI, Yi X, Schuetze DP, Pal R et al. Evidence for an unanticipated relationship between undifferentiated pleomorphic sarcoma and embryonal rhabdomyosarcoma. Cancer cell 2011; 19: 177–191.

21 Rudzinski ER, Anderson JR, Chi YY, Gastier-Foster JM, Astbury C, Barr FG et al. Histology, fusion status, and outcome in metastatic rhabdomyosarcoma: A report from the Children’s Oncology Group. Pediatric blood & cancer 2017; 64.

22 Shern JF, Chen L, Chmielecki J, Wei JS, Patidar R, Rosenberg M et al. Comprehensive genomic analysis of rhabdomyosarcoma reveals a landscape of alterations affecting a common genetic axis in fusion-positive and fusion-negative tumors. Cancer Discov 2014; 4: 216–231.

23 Shukla N, Ameur N, Yilmaz I, Nafa K, Lau CY, Marchetti A et al. Oncogene Mutation Profiling of Pediatric Solid Tumors Reveals Significant Subsets of Embryonal Rhabdomyosarcoma and Neuroblastoma with Mutated Genes in Growth Signaling Pathways. Clin Cancer Res 2012.

24 Smith LM, Anderson JR, Coffin CM. Cytodifferentiation and clinical outcome after chemotherapy and radiation therapy for rhabdomyosarcoma (RMS). Medical and pediatric oncology 2002; 38: 398–404.

25 Sokolowski E, Turina CB, Kikuchi K, Langenau DM, Keller C. Proof-of-concept rare cancers in drug development: the case for rhabdomyosarcoma. Oncogene 2014; 33: 1877–1889.

26 Sriraman SK, Geraldo V, Luther E, Degterev A, Torchilin V. Cytotoxicity of PEGylated liposomes co-loaded with novel pro-apoptotic drug NCL-240 and the MEK inhibitor cobimetinib against colon carcinoma in vitro. Journal of controlled release: official journal of the Controlled Release Society 2015; 220: 160–168.

27 Svalina MN, Keller C. YAPping about differentiation therapy in muscle cancer. Cancer cell 2014; 26: 154–155.

28 The Broad Institute of MIT and Harvard. 2019.

29 Thompson MC, Fuller C, Hogg TL, Dalton J, Finkelstein D, Lau CC et al. Genomics identifies medulloblastoma subgroups that are enriched for specific genetic alterations. J Clin Oncol 2006; 24: 1924–1931.

30 Wachtel M, Dettling M, Koscielniak E, Stegmaier S, Treuner J, Simon-Klingenstein K et al. Gene expression signatures identify rhabdomyosarcoma subtypes and detect a novel t(2;2)(q35;p23) translocation fusing PAX3 to NCOA1. Cancer Res 2004; 64: 5539–5545.

31 Weis SL, Goheer MA, Helms WS. NDA 206,192: Pharmacology/Toxicology NDA Review and Evaluation, 2015.

32 Yamaguchi T, Kakefuda R, Tajima N, Sowa Y, Sakai T. Antitumor activities of JTP-74057 (GSK1120212), a novel MEK1/2 inhibitor, on colorectal cancer cell lines in vitro and in vivo. International journal of oncology 2011; 39: 23–31.

33 Yohe ME, Gryder BE, Shern JF, Song YK, Chou HC, Sindiri S et al. MEK inhibition induces MYOG and remodels super-enhancers in RAS-driven rhabdomyosarcoma. Science translational medicine 2018; 10.

